# Scaffold-free 3D-Cell Co-Culture Model System for the Study of Metastatic Cancer Cell Behavior in the Brain TME

**DOI:** 10.1101/2025.10.31.685975

**Authors:** Pratistha Sarkar, Shreya Ahuja, Iulia M. Lazar

## Abstract

Cancer is a complex disease involving dynamic interactions between cancer, stromal, and infiltrating immune cells, as well as between these cells and the extracellular matrix components of the tumor microenvironment. Brain metastases arise primarily from solid tumors and often result in fatal outcomes. An in-depth understanding of the complex intercellular interactions that evolve in the brain microenvironment is essential to enabling early cancer diagnostics and improving patient outcomes. The protected tumor microenvironment of the brain hinders, however, direct access, impeding the execution of mechanistic studies and limiting the ability to derive meaningful insights. Several *in vitro* 2D and 3D model systems have been developed to circumvent this problem, none, however, without limitations. The 2D models fail to recapitulate the 3D architecture of the *in-vivo* environment lacking therefore physiological relevance, while the 3D models present challenges related to the lack of control over cell positioning, lack of vascularization, contamination from non-human scaffolds, batch-to-batch reproducibility, and high production costs. To overcome some of these limitations, we developed an *in vitro* scaffold-free 3D tumor model system to simulate the *in vivo* brain metastatic niche. The model was constructed from human brain endothelial cells (HBEC-5i) and two different cancer cell lines derived from breast (MDA-MB-231/triple negative and SK-BR-3/HER2+) and aggressive ovarian (SK-OV-3) cancers. The development of the model relies on a newly identified affinity between the endothelial and cancer cells that enables them to self-assemble in 3D networked constructs, a feature facilitated by the high collagen production by endothelial cells and the secretion of key chemokines by both endothelial and cancer cells. The model mimics the attachment of metastasized cancer cells to the brain microvasculature, enabling the study of temporal changes in endothelial morphology and molecular signaling processes that sustain cancer cell migration, survival, proliferation, and angiogenic processes. Moreover, the model exhibits long-term stability, reproducibility, and effectiveness in evaluating anti-cancer agents. Altogether, the scaffold-free, simple 3D *in vitro* model systems provides a low cost, physiologically relevant tool for studying the dynamic molecular crosstalk between cancer and brain endothelial cells, and for investigating the fundamental biological processes that unfold in the tumor microenvironment.

## Introduction

Endothelial cells form the inner layer of blood vessels and have primary functions in controlling vascular tone and blood flow, mediating inflammation, promoting angiogenesis, and supplying tissues with oxygen and nutrients (Ghitescu & Robert, 2002; Krüger-Genge et al., 2019; Trimm & Red-Horse, 2023). Depending on the tissue of origin and vascular bed, endothelial cells display a diverse range of morphological and biochemical signaling pathways, gene expression patterns, and membrane protein profiles. Specifically, the brain endothelial cells are different from the endothelial cells of other organs by exhibiting (i) higher levels of tight junction proteins (claudins 5-CLDN5 and occludin-OCLN) that are essential for the formation of the blood brain barrier (BBB), (ii) increased mitochondrial content for supporting a high metabolic demand, (iii) reduced pinocytic activity to minimize the entry of non-specific substances form the blood into the brain, and (iv) elevated levels of intake transporter proteins (e.g., the glucose transporter 1-GLUT1/SLC2A1, organic anions and thyroid hormones solute carrier SLCO1C1, and long chain fatty acids sodium dependent lysophosphatidylcholine symporter 1 NLS1/ MFSD2A) as well as efflux pumps for removing various xenobiotics out of the cells (e.g., the ATP-Binding Cassette Transporter G2 ABCG2, a multidrug resistance protein) (Arvanitis et al., 2020; Krüger-Genge et al., 2019). Differences in endothelial cell-membrane post-translational modifications such as glycosylation have been also reported (Ghitescu & Robert, 2002; Krüger-Genge et al., 2019). In particular, the role of the luminal cell-membrane anchored proteoglycans, glycoproteins and carbohydrates (the glycocalyx) in signaling, inflammation, vascular permeability, mechanotransduction, and pathophysiology, has been extensively documented (Moore et al., 2021; Reitsma et al., 2007).

The endothelial cells are also one of the major stromal cells of the tumor microenvironment (TME). Signaling processes between the tumor and TME cells play a crucial role not only in stimulating tumor growth, migration and metastatic behavior, but also in remodeling the behavior of endothelial cells and angiogenic programs. The tumor cells can target the endothelial cells in various ways. Via intercellular communication, tumor and tumor-associated stromal cells secrete various soluble pro-angiogenic proteins, importantly VEGF, FGF2, and PDGF that activate the tyrosine kinase receptors on endothelial cells to further upregulate downstream angiogenesis pathways (Z.-L. Liu et al., 2023). Through connexin-43 dependent gap junctions, tumor cells exchange ions and small metabolites with endothelial cells, which can support then tumor progression and metastasis (Wu & Wang, 2019). Furthermore, by releasing exosomes containing pro-angiogenic proteins, mRNAs and microRNAs, tumor cells trigger angiogenic gene expression in endothelial cells to support their own growth and metastasis (Olejarz et al., 2020). Indirectly, the tumor cells modulate the behavior of other stromal cells and the microenvironment by altering the physiological conditions, including the pH, oxygen levels, temperature, and TME composition, to induce endothelial cell responses (Lopes-Bastos, 2016). For example, as a result of the Warburg effect, cancer cells produce lactate which is released in the TME via the monocarboxylate transporter (MCT-4). When imported by endothelial cells through the MCT-1 transporter, lactate activates the NF-kB/IL-8 pathway. IL-8 produced through this mechanism acts in an autocrine manner to activate endothelial cells and stimulate angiogenesis (Lidonnici et al., 2022).

To investigate the intricate interactions between cancer and endothelial cells, tumor intravasation, angiogenic or immune signaling, or drug screening and delivery, several model systems have been developed (Chen et al., 2018; Hachey et al., 2023; Lopez-Vince et al., 2024; Oh et al., 2020; Ye et al., 2024; Zervantonakis et al., 2012). *In vitro*, two-dimensional (2D) cancer/endothelial co-culture models have been extensively explored. Such 2D models are relatively cost-effective, easy to generate and use, and can provide valuable insights into cell morphology, the dynamics of fundamental biological processes, and pharmacokinetic behavior. Nonetheless, such models fail to recapitulate the intricate three-dimensional (3D) architecture of the *in-vivo* cellular environment. To address this shortcoming, alternative 3D-spheroid and organoid models that simulate tumor morphology, extracellular matrix (ECM) composition, the effect of chemical gradients (cytokines, chemokines, and other soluble factors), and cell-cell and cell-matrix interactions have been developed (Abuwatfa et al., 2024; Peng et al., 2025; Sharma et al., 2024). To provide structural support and imitate the ECM, the 3D co-culture models are typically constructed on a scaffold prepared from various natural or synthetic materials including decellularized tissues, polymers, hydrogels, or hybrid materials (Abuwatfa et al., 2024; Brassard-Jollive et al., 2020; Hippler et al., 2019; Sharma et al., 2024). Among these, animal-derived matrices such as the ECM hydrogel derived from mouse sarcoma (Matrigel) and the rat tail collagen I are commonly used due to their widespread availability (Andrée et al., 2019). To further increase biological relevance, such cancer/endothelial co-culture models have been also integrated in microfluidic devices, which enable the investigation of physiological flow conditions and shear stress effects on cellular behavior (Chen et al., 2018; Hachey et al., 2023; Zervantonakis et al., 2012). In one such example, the microfluidic setup was used to quantify the spatial and temporal dynamics of tumor/vasculature/ECM interactions and angiogenic sprouting specific to inflammatory breast cancer (Gadde et al., 2020). These model systems raise, however, concerns about the potential impact of contaminants of non-human origin, immunogenicity, and poor reproducibility due to variations in the composition of the scaffolds. Additionally, due to increased complexity and heterogeneity of the culture environment, the 3D systems are more difficult to reproduce than the 2D models (Abuwatfa et al., 2024). A scaffold-free strategy relying on the ability of cells to self-organize without needing synthetic or animal-derived matrices would address these issues, and would be an ideal tool for representing the *in-vivo* tumor biology and performing high-throughput drug screening and cancer modeling (Mason & Öhlund, 2023).

For simulating the *in-vivo* tumor microenvironment and studying angiogenic processes and vasculature biology, most existing endothelial models utilize macrovasculature human umbilical vein endothelial cells (HUVEC). Due to ease in accessibility, isolation, and culture, HUVEC cells have been even utilized for developing *in vitro* microfluidic models of the BBB network, despite lacking many properties of the brain microvasculature endothelial cells (Bang et al., 2017). A study by Eigenmann *et al*. has shown, however, that human brain microvasculature endothelial cells (HBMEC) that form a more restrictive barrier with a more favorable trans endothelial electrical resistance (TEER) and lower permeability, represents a more suitable cell line for modeling the BBB and performing high-throughput permeability screening (Eigenmann et al., 2013). Another study demonstrated that the immortalized human cerebral microvascular endothelial cell line (HCMEC/D3) isolated from the temporal lobe, and expressing platelet endothelial cell adhesion molecule 1 (PECAM-1), junctional adhesion molecule-A (JAM-A), VE-cadherin (CDH5), and tight junction CLDN3/CLDN5 and OCLN, is suitable for developing physiologically relevant *in vitro* 3D BBB models for studying barrier function and regulation, signaling mechanisms, and screening for drug uptake and neurotoxicity (Helms et al., 2016). As an alternative to HCMEC/D3 cells, the cerebral cortex microvascular endothelial cell line (HBEC-5i) was used by Bolden *et. al*., due to its high expression of tight junction proteins, high TEER, and low permeability when compared to other immortalized BMECs (Bolden et al., 2023). In addition to the above-described immortalized brain endothelial cells, primary or low passage BMECs have been also utilized for building *in vitro* BBB models. The limited culture period can, however, compromise the barrier maturity and TEER stability. Human pluripotent stem cells (hPSC) represent another resource for producing endothelial cells, notwithstanding their limitations related to incomplete maturation, heterogeneous cellular identity, early senescence, and poor *in vivo* mimicry (He et al., 2014).

Metastasized cancer cells that cross the BBB co-opt the brain microvessel capillaries to draw oxygen and nutrients, and to support their own survival and growth while they adapt their metabolism to the new tumor microenvironment (McDonald et al., 2023; Srinivasan et al., 2021). In this manuscript we describe the development of a novel, proof-of-concept, scaffold-free 3D endothelial/cancer cell model system aimed at mimicking the brain metastatic niche. The model leverages two novel findings in our laboratory, i.e.: (i) Preliminary work has identified an affinity between brain endothelial and cancer cells that enables them to self-assemble in 3D structures in a scaffold-free environment, and mimic the attachment of metastasized cancer cells to the brain microvasculature; and (ii) The proteomic profiling of brain and cancer cell secretomes revealed that the endothelial cells secrete considerably more collagen into their environment than other cell types. The model was built with HBEC-5i cells which present multiple advantages over other immortalized brain endothelial cells due to their high expression of proteins that delineate key interactions with cancer cells, and three distinct cancer cell lines of two different tissues of origin, i.e., MDA-MB-231 (triple negative/TN, lacking estrogen, progesterone and HER2 receptors) and SK-BR-3 (HER2+) - two cancer cell lines representing frequently brain metastasized breast cancers, and SK-OV-3 - an aggressive epithelial ovarian cancer cell line representing a cancer with poor prognosis that seldom metastasizes to the brain, limiting thereby the opportunities for its study. Our scaffold-free co-culture model exhibits a 3D networked structure, direct cell-to-cell contact, cancer cell-dependent changes in endothelial cell morphology, and long-term stability, establishing a valuable tool for investigating the dynamic molecular crosstalk between endothelial and cancer cells. We envision that the platform will find broad utility for investigating survival, angiogenic, adhesion, and migratory programs in the TME for performing drug screening, and for developing immunotherapeutic approaches.

## Methods

### Reagents

Stable turboFP602 transfected MDA-MB-231 and SK-BR-3 breast and turboGFP transfected SK-OV-3 human ovarian cancer cell lines were purchased from Cells Online LLC (Milpitas, CA). HBEC-5i human brain endothelial cells were obtained from ATCC (Manassas, VA). Cell culture media [RPMI 1640 (1X), McCoy’s 5A (modified), DMEM high glucose (HG)], fetal bovine serum (FBS), GlutaMAX, Pen-Strep, puromycin, G418, Dulbecco’s phosphate buffered saline (DPBS), trypsin (0.25 %)/EDTA (0.53 mM), and propidium iodide (PI) dead cell stain were purchased from ThermoFisher (Carlsbad, CA). Lapatinib ditosylate was acquired from Selleck (Houston, TX), and endothelial cell growth supplement (ECGS) from Sigma Aldrich (St. Louis, MO). Primary ZO1 rabbit IgG mAb and secondary anti-rabbit IgG antibody conjugated to AlexaFluor 488 were purchased from Cell Signaling Technology (Danvers, MA), and primary Vimentin mouse IgG mAb and secondary anti-mouse IgG antibody conjugated to CruzFluor 488 from Santa Cruz Biotechnology (Dallas, TX). ActinRed 555 ReadyProbe Rhodamine phalloidin F-actin stain was from ThermoFisher, and DAPI nuclear counter stain from Cell Signaling Technology. EasyProbe green 488 dead cell stain was purchased from ABP Biosciences (Rockville, MD). PermaCell 24 well plates with cell culture inserts with 8 µm pore polycarbonate membranes and 8-well chamber slides were ordered from MatTek Life Sciences (Ashland, MA). Pipettes and Nunc EasYFlask polystyrene cell culture flasks with Nunclon Delta surface treatment were obtained from Life Technologies (Carlsbad, CA).

### Cell culture

For stock preparation, the cells were initially cultured in the manufacturer’s recommended growth medium, i.e., turboGFP SK-OV-3 and turboFP602 SK-BR-3 in McCoy’s 5A supplemented with FBS (10-15 %), glutamine (2 mM), and G418 (250 µg/mL); turboFP602 MDA-MB-231 in RPMI 1640 (1X) supplemented with FBS (10 %) and puromycin (10 µg/mL); and, HBEC-5i in DMEM/HG supplemented with FBS (∼15 %), ECGS (∼27 µg/mL), and Pen-Strep (0.5 %). For downstream experiments, the cells were grown in DMEM/HG supplemented with FBS (10 %) and Pen-Strep (0.5 %). All concentrations represent the final concentrations in the cell culture medium, with liquid solution percentages representing v/v concentrations. The cell cultures were kept in a humidified incubator at 37 °C with 5 % CO_2_, and the culture medium was changed every other day.

### Cell co-cultures

Two distinct methods were used for developing the scaffold-free co-culture model of cancer and endothelial cells. All experiments were performed in duplicate. In the first approach, the endothelial cells were mixed with either green fluorescent SK-OV-3 or red fluorescent MDA-MB-231 cells in T25 cm^2^ Nunc flasks in a ratio of 9:1 (i.e., 180,000 HBEC-5i: 20,000 cancer cells), or a ratio of 1:1 at double density seeding (i.e., 200,000 of each HBEC-5i and cancer cells). The co-culture mixtures were maintained in DMEM/HG medium supplemented with ECGS (∼27 µg/mL), FBS (10 %), and Pen-Strep (0.5 %). For HBEC-5i/MDA-MB-231 co-cultures, ECGS was added to the culture medium only until day 16. In the second approach, the HBEC-5i endothelial cells were seeded at low confluence (200,000 cells) in a T25 cm^2^ Nunc flask and maintained in DMEM/HG medium supplemented with ECGS (∼27 µg/mL), FBS (10 %), and Pen-Strep (0.5 %) for ∼10 days until 3D networked structures developed. At that stage, 200,000 cancer cells were added to the flask.

### Fluorescence microscopy

An inverted Eclipse TE2000-U epi-fluorescence microscope (Nikon Instruments Inc., Melville, NY, USA), with Plan Fl l0X DL PhObj NA 0.30 WD 16.0mm PH1 and Plan Fl ELWD DM Phase 20X NA 0.45 WD 7mm-8.lmm CC objectives, was used to visualize the cancer-endothelial co-culture model systems. The excitation/emission maxima for green fluorescent SK-OV-3 cells and red fluorescent MDA-MB-231/SK-BR-3 cells were 482/502 nm and 574/602 nm, respectively. An AURA III solid state triggerable light source (Lumencor, Inc. Beaverton, OR) using a DAPI/FITC/TRITC/CY5 Quad filter (Chroma Technology Corp., Bellows Falls, VT) was used to illuminated the cells. The exposure time was 100 ms. An ORCA-Flash 4.0 LT+sCMOS digital camera (Hamamatsu Photonics, Bridgewater township, NJ) and the Nikon’s NIS-Elements Advanced Research imaging platform, version 5.11.01, were utilized for acquiring and processing the images. For immunofluorescence staining, the cells were cultured in chamber slides, fixed with chilled methanol (-20°C) for 15 min, and then exposed to blocking buffer [PBS + BSA (5 %) + Triton X-100 (0.3 %)] for 1 h at room temperature, primary antibody overnight at 4°C, and secondary antibody for 1.5 h at room temperature in the dark. DAPI was added to all chambers and the cells were incubated for 5 min. The antibodies were prepared in a PBS solution supplemented with BSA (1 %) and Triton X-100 (0.3 %) in the following concentrations: primary ZO1 rabbit IgG (0.07 µg/mL), secondary anti-rabbit IgG-AlexaFluor 488 (2 µg/mL), primary vimentin mouse IgG (2 µg/mL), and secondary anti-mouse IgG-CruzFluor 488 (1 µg/mL). ActinRed 555 (2 drops/mL) and DAPI (1 µg/mL) staining was performed after completing the secondary antibody staining procedure.

### Transwell migration assay

Transmigration assays of green fluorescent SK-OV-3 and red fluorescent SK-BR-3 cells were performed in transwell plates with polycarbonate membrane inserts of 8 µm pore size. HBEC-5i cells (50,000/well) were seeded in the bottom chamber in 600 µL medium [DMEM/HG, FBS (10 %), ECGS (∼27 µg/mL), Pen-Strep (0.5 %)] and incubated for 24 h in a water-jacketed incubator at 37 °C with 5 % CO_2_. For the control wells, the bottom chamber contained only the cell culture medium, but no HBEC-5i cells. After 24 h, when the HBEC-5i cells reached ∼60 % confluence, cancer cells suspended in 100 µL DMEM/HG supplemented with FBS (10 %) and Pen-Strep (0.5 %) were added to the upper inserts (∼50,000 cells/insert) for both experimental and control wells, and the transwell plates were incubated for up to 5 days in an incubator. The migration of fluorescent cancer cells from the inserts toward the endothelial cells in the bottom chambers was monitored by fluorescence microscopy. After 5 days of incubation, the inserts were removed and images were taken from both control and experimental bottom wells.

### Drug treatment

HBEC-5i and red fluorescent SK-BR-3 cells were mixed in a proportion of 1:1 and co-cultured in DMEM/HG medium supplemented with FBS (10 %), ECGS (∼27 µg/mL), and Pen-Strep (0.5 %) for up to 5 days, when Lapatinib drug (1 µM or 10 µM) was added to the co-culture system. HBEC-5i and SK-BR-3 monocultures were also incubated separately with Lapatinib (10 µM). All drug treatments were performed for up to 72 h. The experiments were performed in duplicate and the dead cells were visualized with EasyProbe green dead cell stain.

### Live/dead cell assay

The long-term viability of cells was monitored by visualizing the presence of dead cells, up to 28 days in co-culture, by fluorescence microscopy. Propidium iodide was used for visualizing dead green fluorescent SK-OV-3 cells (excitation/emission maxima 535/617 nm when bound to dsDNA; 5 µg/mL in the culture medium), and EasyProbe green dead cell stain for visualizing dead red fluorescent MDA-MB-231 and SK-BR-3 cells (excitation/emission maxima 500/530 nm; 2 drops/ml in the culture medium). The cells were incubated for 30 min with the dead cell stains in the culture medium in a 37 °C humidified incubator with 5 % CO_2_, and visualized by fluorescence microscopy directly in the culture medium without any further preparation.

### Proteomic data analysis

A detailed description of proteomic sample generation and data acquisition and processing methodology is provided in previous work (Ahuja & Lazar, 2021; Karcini & Lazar, 2022). Briefly, the cells were cultured in DMEM/HG supplemented with FBS (10 %) and Pen-Strep (0.5 %). Cell-membrane proteins were isolated from cultures produced in serum-rich and serum-free media (Karcini & Lazar, 2022), while cell secretomes were collected from the supernatant of serum-starved cells for 48 h that were concentrated with Amicon Ultra-15 centrifugal filters (3000 MWCO). After proteolytic digestion with trypsin and sample cleanup, data-dependent mass spectrometry proteomic data were acquired with a nano-LC (EASY-nLC 1200) separation system interfaced to a hybrid quadrupole-Orbitrap^TM^ mass spectrometer (Q Exactive^TM^, ThermoFisher Scientific). RAW MS files were processed with the Proteome Discoverer (v.2.5) software package by using the Sequest HT search engine and a minimally redundant/reviewed *Homo sapiens* database (August 30, 2022 download, 20399 entries). Peptide target false discovery rate thresholds were set to 0.01/0.03 (stringent/relaxed). Semiquantitative comparisons of cell-membrane and secretome data were performed based on peptide spectrum match (PSM) counts, after normalizing the data from each cell line based on total PSMs per dataset. Cord diagrams from the proteomic data were created with RAWGraphs tools (https://rawgraphs.io).

## Results

### Development of the scaffold-free 3D endothelial/cancer cell co-culture model

Focus was placed on developing a system in which co-evolving cancer and endothelial cells form assemblies that mimic the brain microvasculature which provides cancer cells with nutrients and oxygen, facilitating tumor progression and metastasis. HBEC-5i endothelial cells were chosen due to their ability to form dense 2D monolayer networks with well-developed functional tight junctions that accurately replicate the *in-vivo* brain microvasculature. The HBEC-5i cells also exhibit high-abundance expression of key proteins that are critical to the establishment of cancer cells in the TME, such as adhesion (CDH5, VCAM1/CD106, ICAM1/CD54), transport (ABC transporters), and tight junction (CLDN5, OCLN) proteins, as well as receptors critical to modulating immune and inflammatory responses (TNF receptor CD40) (*ATCC*, n.d.). Co-culture with astrocytes, which we intend to incorporate in future work, has been shown to increase, or alter, the expression of adhesion/tight junction and ABC transporter proteins (Puech et al., 2018; Sun & He, 2025). The three cancer cell lines that were investigated represent useful preclinical models that are often used to study the fundamental mechanisms of cancer biology, the efficacy of therapeutic drugs, and the mechanisms of drug resistance. Breast cancer MDA-MB-231 (TN) and ovarian cancer SK-OV-3 cells were used to develop the model, and breast cancer SK-BR-3 (HER2+) to test the response to exposure to a small-molecule receptor Tyr kinase (RTK) inhibitor, Lapatinib.

At low confluence, in monoculture, the HBEC-5i cells presented a spindle-shaped morphology. With increased confluence, typically after 10 days of culture, 3D tubular-like structures were spontaneously formed in regular culture medium, without any need for scaffolding support (**Figures 1A-C**). Fluorescence microscopy imaging revealed a healthy endothelial culture capable of forming tight junctions, a key indicator that the cells maintained their original functionality. This was evidenced by visualizing cell-membrane localized ZO-1 proteins, cortical ring actin, and cytoplasmic vimentin with a characteristic denser accumulation in the perinuclear region (**Figures 1D-F**). The ability to develop networked/tubular-like structures is a known feature of endothelial cells grown in isolation (Andrée et al., 2019). When grown in co-culture, however, the endothelial and cancer cells displayed a unique self-assembly behavior that has been overlooked in previous studies. The use of transfected red-fluorescent breast (tFP602 red) and green-fluorescent ovarian cancer (tGFP green) cells enabled the observation of a remarkable behavior of cells, i.e., the cancer and endothelial cells self-assembled in 3D constructs that mimicked the topology of brain microvasculature (∼10 µm diameter, ∼40-200 µm length (Ji et al., 2021)) where the cancer cells attach and thrive until bursting in a new tumor. Proteomic profiling studies in our laboratory also revealed that the HBEC-5i cells secrete a significant amount of collagen compared to the cancer cells, which we believe served as a natural scaffolding material that supported the necessary cell-cell interactions that led to the formation of microvascular-like structures.

**Figure 1.**
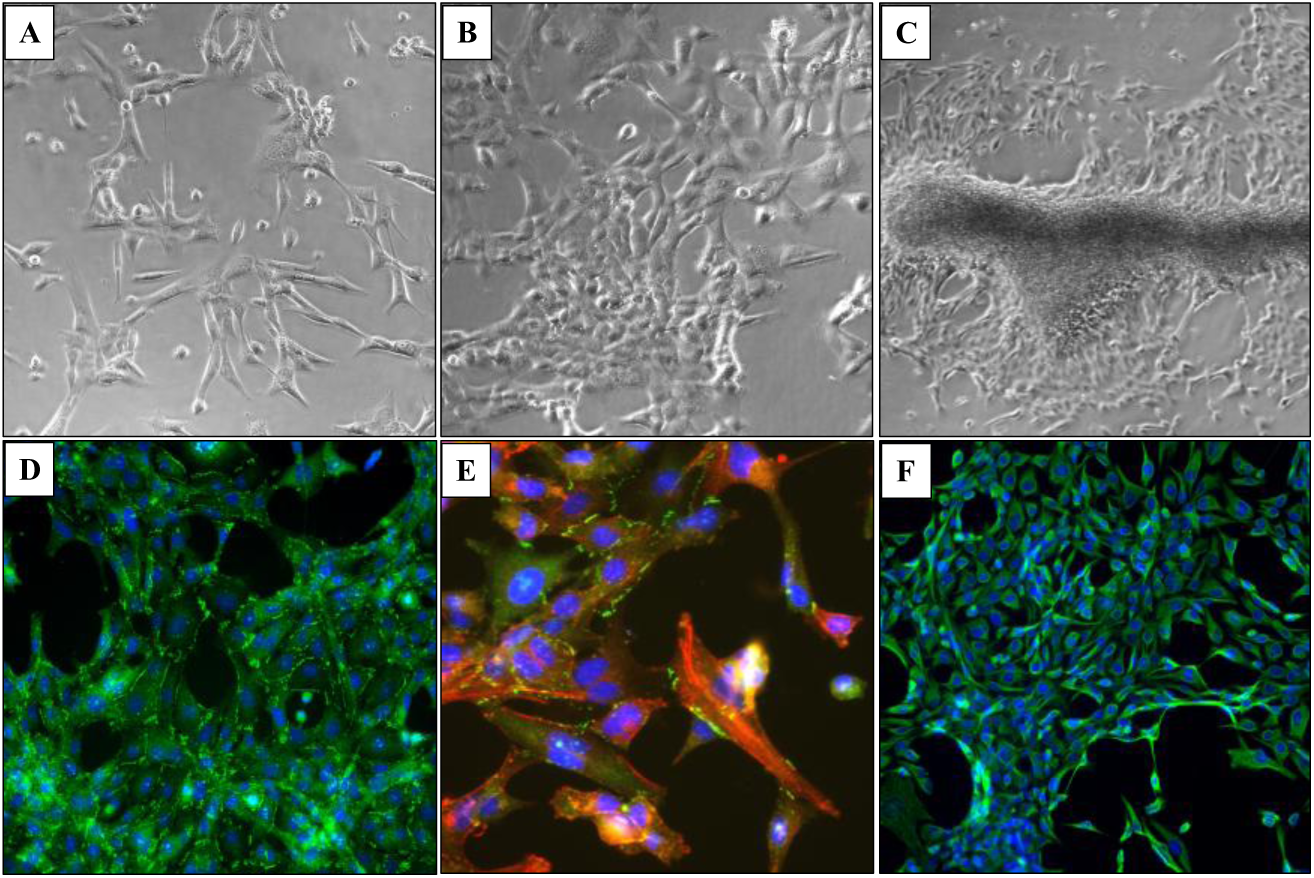
Microscopy images of proliferating HBEC-5i cells. (**A**) Endothelial cells at low density seeding; (**B**) Multi-cellular assemblies of endothelial cells formed at higher confluence; (**C**) Self-organization of endothelial cells in 3D aggregates that mimic the tubular microvasculature; (**D**) Immunofluorescent visualization of ZO-1 accumulation at nascent endothelial cell contact sites (green dots and lines); (**E**) Immunofluorescent staining of ZO-1 (green), cytoplasmic actin (red), and cell nuclei (blue); (**F**) Immunofluorescent staining of vimentin (green) and cell nuclei (blue).

Based on these findings, we developed two approaches for building 3D endothelial/cancer co-culture models in which the endothelial and cancer cells were either pre-mixed and allowed to co-evolve, or the cancer cells were added to pre-formed endothelial constructs. For the 1^st^ approach, the HBEC-5i and MDA-MB-231 or SK-OV-3 cancer cells were mixed in a proportion of 9:1 or 1:1 to mimic either early or later stages of metastatic growth, respectively (**Figures 2** and **3**). For the 2^nd^ approach, HBEC-5i cells were seeded in a flask and allowed to evolve for ∼10 days until cellular networks of microvascular-like structures formed, time at which the cancer cells were added (**Figure 4**). The co-culture was then maintained using the same medium and cell culture conditions.

**Figure 2.**
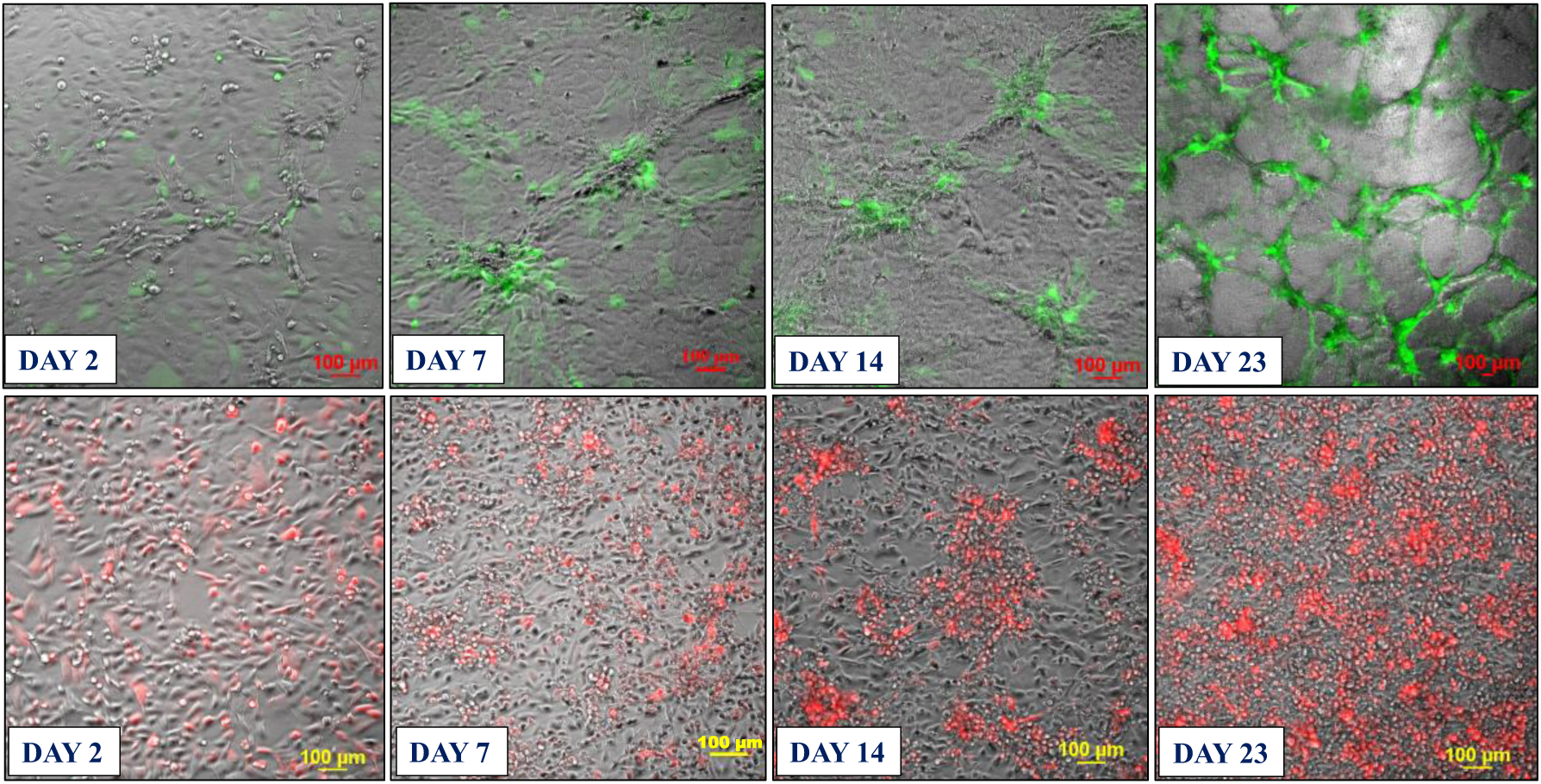
Development of the co-culture model system prepared from premixed HBEC-5i and stable transfected fluorescent cancer cells (1:1 mix). **Top panel**: HBEC-5i/SK-OV-3 (turbo GFP/green) model; **Bottom panel**: HBEC-5i/MDA-MB-231 (turbo FP602/red) model. Cells were seeded in equal proportion (200,000 each, ∼30 % confluence) in a 25 cm^2^ culture flask and allowed to evolve for ∼30 days. Self-organization of endothelial cells in 3D aggregates and accumulation of cancer cells around these structures was observable after ∼7 days (80-90 % confluence). After ∼21 days, clearly delineated 3D networked constructs incorporating the vast majority of cancer cells were observable.

**Figure 3.**
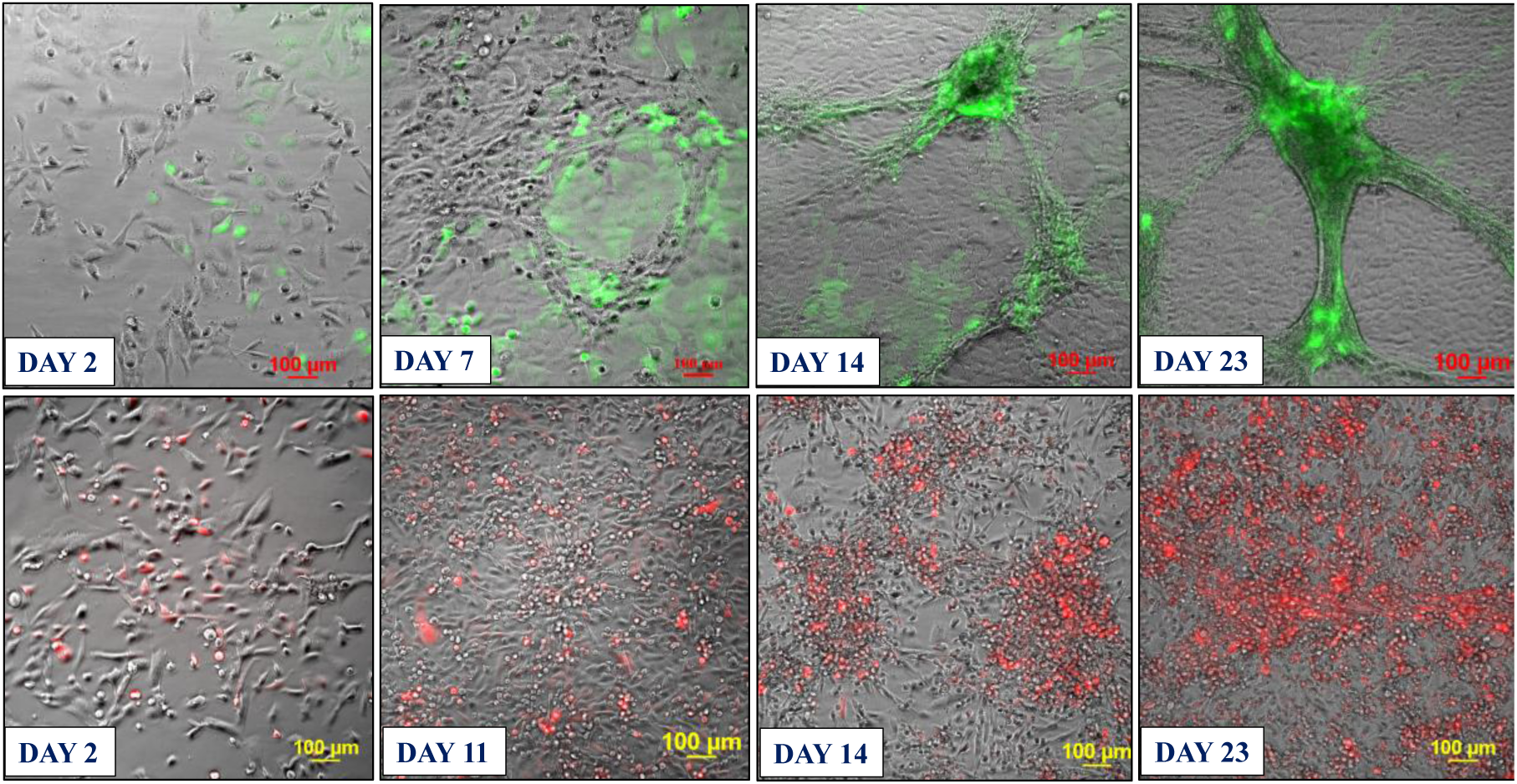
Co-culture model system prepared from premixed HBEC-5i and stable transfected fluorescent cancer cells (9:1 mix). **Top panel**: HBEC-5i/SK-OV-3 (turbo GFP/green) model; **Bottom panel**: HBEC-5i/MDA-MB-231 (turbo FP602/red) model. Cells were seeded in a proportion of HBEC-5i (180,000 cells)/cancer cells (∼20,000) in a 25 cm^2^ culture flask and allowed to evolve for ∼30 days. Self-organization of endothelial cells in 3D aggregates and accumulation of cancer cells in these structures was observable after ∼7 days. After ∼21 days, clearly delineated 3D networked constructs were observable.

**Figure 4.**
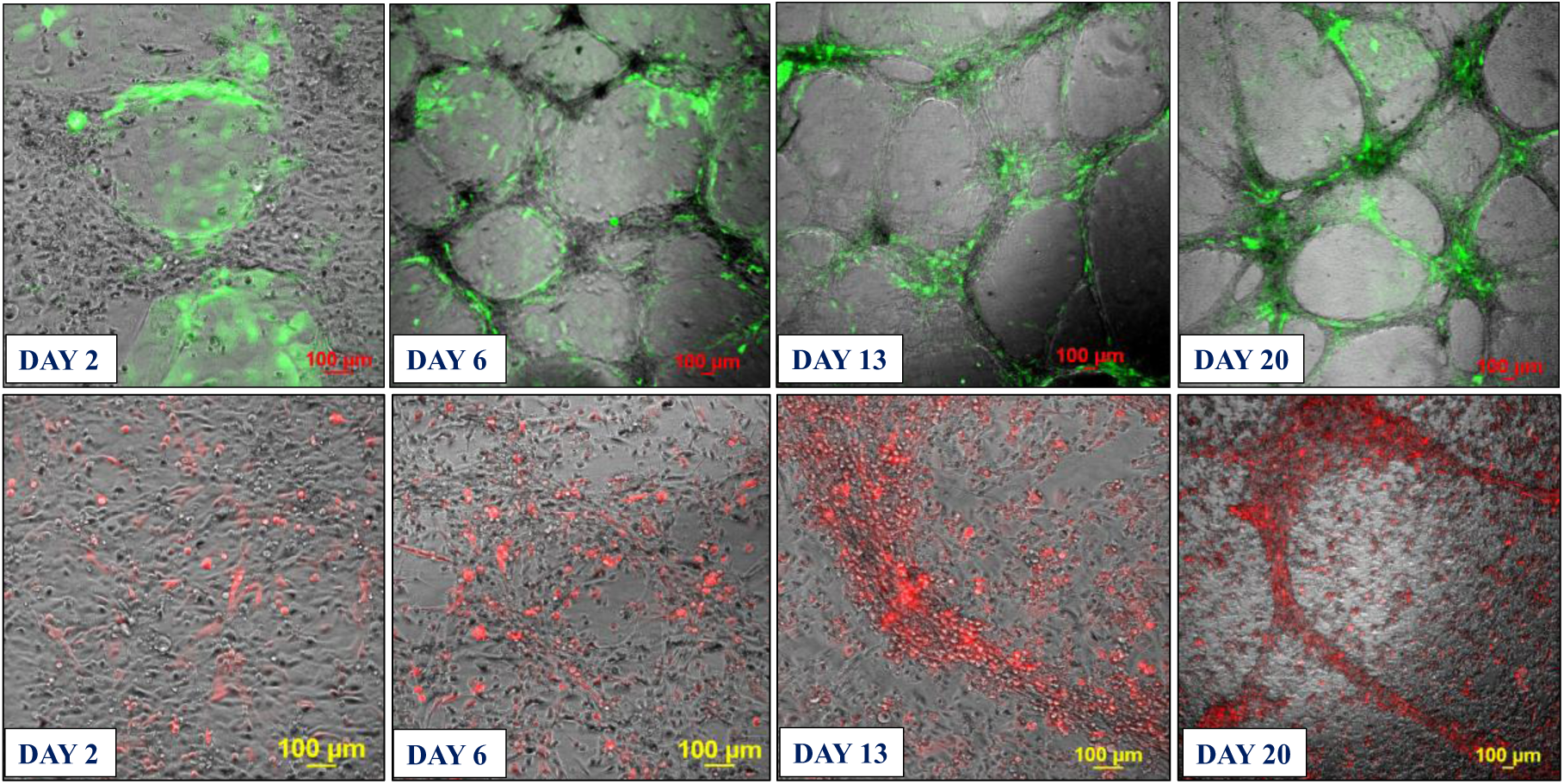
Co-culture model system prepared from pre-formed HBEC-5i networked structures to which stable transfected fluorescent cancer cells were added. **Top panel**: HBEC-5i/SK-OV-3 (turbo GFP/green) model; **Bottom panel**: HBEC-5i/MDA-MB-231 (turbo FP602/red) model. HBEC-5i cells (∼200,000) were seeded in a 25 cm^2^ culture flask and allowed to evolve until they formed 3D networked structures (∼10 days), time when ∼200,000 cancer cells were added to the flask. The migration of SK-OV-3 and MDA-MB-231 cells toward HBEC-5i structures was observable already after 2 days of co-culture. The evolution of endothelial/cancer constructs was monitored for an additional 3 weeks.

### Characterization of the endothelial/cancer cell constructs

Whether added from start or after the endothelial structures were formed, the cancer cells migrated toward, attached themselves, influenced the morphology, and continued to grow with/within the endothelial constructs. The endothelial/cancer assemblies formed on un-coated plastic surfaces, without the addition of contaminating animal or synthetic ECM/lamina or scaffold components, and were stable for >4 weeks. The final constructs had consistent morphology in replicate preparations and demonstrated uniform growth, long-term stability, and reproducible behavior, leading to progressively better delineated fluorescent cancer cell accumulations as the co-cultures evolved from day 2 to 23 (**Figures 2-4**).

The morphology of constructs that evolved in co-culture varied, however, from one cancer cell line to another. The HBEC-5i/SK-OV-3 co-cultures generated network constructs similar to the ones produced from HBEC-5i alone, while the HBEC-5i/breast cancer co-cultures generated rather discontinuous, spotty networks that formed slower than the HBEC-5i/SK-OV-3 networks. The cancer cells accumulated, however, always in the areas of higher-density endothelial structures, suggesting the presence of paracrine chemokine gradients and inter-cellular interactions that support cancer cell migration and attachment in order to gain survival benefits. The mixing ratio of endothelial-to-cancer cells did not appear to impact the formation of constructs, but the HBEC-5i cells proliferated at a faster rate in the pre-mixed co-cultures, compared to the HBEC-5i monocultures, with prominent structures being already observable after 6-7 days of co-culture vs ∼10 days in monoculture, respectively. The cancer cells stimulated the formation of well-organized 3D endothelial cellular arrangements and network branching points which entirely covered the small flask area within two weeks of culture. Both SK-OV-3 and MDA-MB-231 cells secrete pro-angiogenic factors known to facilitate angiogenic sprouting and vascular tube formation by upregulating the downstream fatty acid-binding protein 4 (FABP4) expression of VEGF signaling in endothelial cells (Elmasri et al., 2009). The observations also align with the findings of a previous study that showed increased microvessel tube formation in HUVEC stimulated by lung cancer cells via upregulated PI3K/AKT and COX2 signaling pathway (Cheng et al., 2017).

In the 2^nd^ approach, within two days of co-culture, the cancer cells were observed to start migrating toward the pre-formed HBEC-5i structures and generate shortly thereafter clearly delineated endothelial/cancer constructs (**Figure 4**). Both cancer cell types exhibited migratory behavior, with SK-OV-3 cells establishing themselves faster in the endothelial networks than the MDA-MB-231 cells. Unlike in the 1^st^ approach when the cells were pre-mixed from start, the continuous network architecture of HBEC-5i/MDA-MB-231 co-cultures was preserved.

The migratory behavior of cancer cells (SK-OV-3 and SK-BR-3) was confirmed with a transwell assay, where HBEC-5i cells were seeded in the bottom wells and the cancer cells in the upper inserts comprising a membrane with 8 µm pores (**Figure 5**). The control wells contained cancer cells in the upper inserts and only culture medium in the bottom wells. After 5 days of incubation, the inserts were removed, and images from the bottom experimental and control wells were acquired. A much larger number of cancer cells were observable in each of the experimental wells that were pre-seeded with HBEC-5i cells, clustering, again, in the areas of higher density endothelial cells, and supporting the observation that the migration of cancer cells in the model system is controlled by chemical cues secreted by the endothelial cells (**Figure 5**).

**Figure 5.**
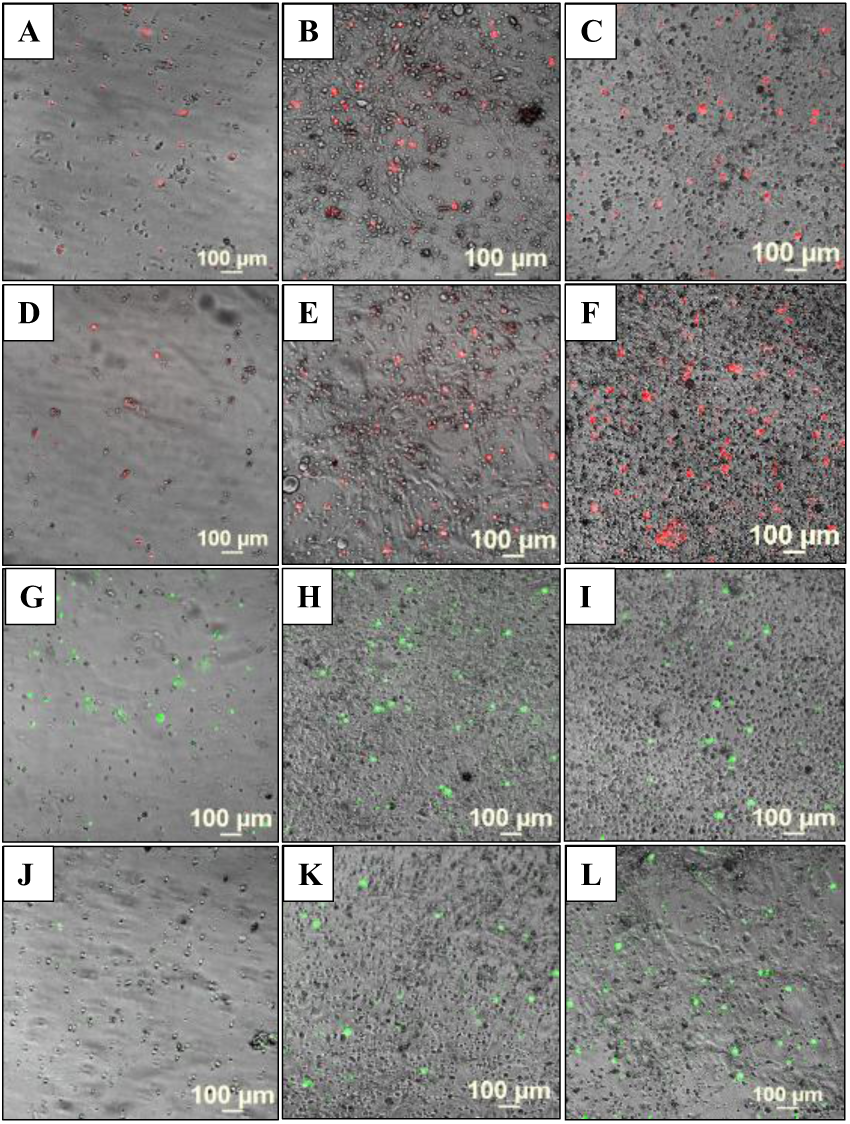
Transwell assay images depicting the migration of cancer cells toward endothelial cells. (**A,D**) Control wells with red fluorescent SK-BR-3 cells seeded in the upper inserts; (**B,C,E,F**) Experimental wells with red fluorescent SK-BR-3 cells seeded in the upper inserts and HBEC-5i in the bottom wells; (**G,J**) Control wells green fluorescent SK-OV-3 cells seeded in the upper inserts; (**H,I,K,L**) Experimental wells with green fluorescent SK-OV-3 cells seeded in the upper inserts and HBEC-5i in the bottom wells. The images were taken from the bottom wells after 5 days incubation.

### Proteomic analysis

Preliminary proteomic profiling of HBEC-5i, SK-OV-3, and MDA-MB-231 cell secretomes and cell-membrane proteins revealed the presence of key cytokines/chemokines, cell adhesion molecules, and growth and angiogenic factors that could support the behavior of cells in the model system (**Figure 6**). While a full analysis of the proteome data is outside the scope of this study, several key findings warrant discussion. As pinpointed above, the endothelial cells secreted several collagens in much higher abundance than the cancer cells, in particular COL1A1, COL1A2, COL6A1, and COL6A3 (see **Supplemental file 1** for a semiquantitative assessment of relative protein abundances in terms of PSMs). These collagens are important for building the ECM, maintaining the integrity of the BBB, and promoting endothelial cell migration and angiogenesis. Moreover, type I collagen has been shown to initiate the formation of 3D capillary networks from proliferating endothelial cells (Whelan & Senger, 2003). The collagen composition of the brain ECM is however highly dynamic, the various members of this superfamily having diverse physiological roles. For example, in the basal lamina of the healthy CNS tissue, the endothelial cells deposit primarily collagen IV, collagens I and VI being rather associated with various diseased states and cerebrovascular pathologies (Gregorio et al., 2018; Wareham et al., 2024). *In vitro*, in cell cultures, the deposition of collagens I and VI could therefore be attributed to an inherent need to establish an ECM foundation that supports inter-cellular signaling and protects against stress. Interestingly, the SK-OV-3 cells also secreted collagens in high abundance, of a different type (COL12A1 and COL18A1), however, which were found in previous studies to be overexpressed in the tumor TME (Q. Zhang et al., 2023). This type of collagen deposition can remodel the ECM and TME composition to promote tumor growth and progression, and in our model served as a scaffolding material that supported the establishment of cell-cell interactions and the formation of 3D constructs. Interestingly, the SK-OV-3 cells in particular, also secreted ECM remodeling matrix metalloproteinases in high abundance (MMP1 and MMP2). These enzymes mediate ECM degradation, enabling cell invasion, and could have been important contributors to the faster formation of HBEC-5i/SK-OV-3 constructs.

**Figure 6.**
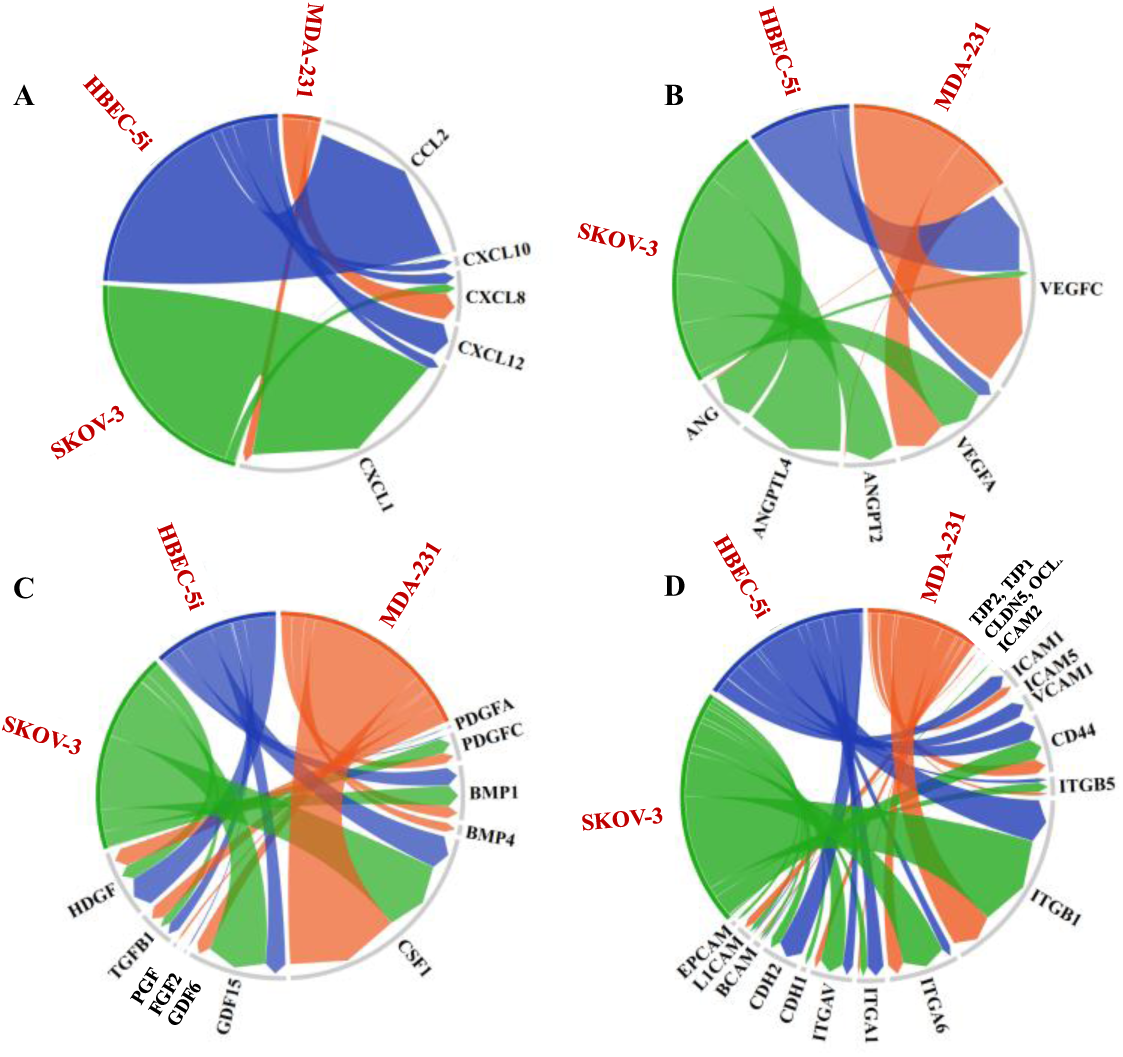
Cord diagrams showing the relative abundance of protein components identified in the cellular secretome or cell-membrane fractions of HBEC-5i, SK-OV-3, and MDA-MB-231 cells, relevant to the formation of 3D endothelial/cancer constructs. (**A**) Chemokines in the cell secretomes (PSM range: 0-200), (**B**) Angiogenic factors in the cell secretomes (PSM range: (0-62), (**C**) Growth factors in the cell secretomes (PSM range: 0-796), (**D**) Adhesion and junction proteins in the cell-membrane proteomes (PSM range: 0-8905).

The cells secreted essential cytokines/chemokines involved in eliciting immune responses and angiogenic processes, modulating the expression of adhesion molecules, and supporting cancer proliferation, EMT, and migration (Salazar & Zabel, 2019; Yoshimura et al., 2023; Zielińska & Katanaev, 2020) (**Figure 6A**). As also shown by our previous microarray profiling studies (Ahuja & Lazar, 2021), the proteomic findings confirmed the presence of CCL2 and CXCL1/8/10/12 in the HBEC-5i secretome. Among these, CCL2 (or MCP-1), an inflammatory cytokine with multiple roles in promoting immune cell infiltration in the brain and tumor cell migration across the endothelium, was present in high abundance. CCL2 was shown to promote breast cancer brain metastasis and the invasion and adhesion of SK-OV-3 cells (Furukawa et al., 2013; Israeli Dangoor et al., 2025). CXCL8 and CXCL1 - secreted by all cells, and CXCL12 - identified only in the HBEC-5i secretome, are known drivers of metastasis by promoting tumor cell motility, adhesion and invasion (Shen et al., 2022, p. 2; Zielińska & Katanaev, 2020). The overexpression of CXCL1 has been shown to increase the proliferation of ovarian cancer cells (Bolitho et al., 2010).

All cells secreted growth and angiogenic factors with key role in stimulating new blood vessel formation and cell growth, including various members of the VEGF, PDGF, FGF, PGH, and ANG family of proteins (**Figure 6B**). VEGF proteins were particularly abundant in all cancer and endothelial cell secretomes, whereas ANG and ANGPT2/ANGPTL4 in SK-OV-3, suggesting a potentially significant role in the formation of the HBEC-5i/SK-OV-3 networked constructs. While ANG (angiogenin) directly stimulates blood vessel formation, ANGPT2 (angiopoietin 2) contributes to vessel destabilization (*GeneCards - Human Genes | Gene Database | Gene Search*, n.d.). Nonetheless, it has been shown that ANGPT2 has a context-dependent role, facilitating endothelial cell proliferation and migration in the presence of VEGF (Gale et al., 2002; Moon et al., 2003). Notable in the secretome of cancer cells was also the high level of CSF1 – a cytokine with role in inflammatory regulatory processes, adhesion and migration (*GeneCards - Human Genes | Gene Database | Gene Search*, n.d.), and of GDF15 – a multifunctional protein involved in many processes of relevance to cancer progression and metastasis such as EMT, remodeling of the TME, immune escape, and development of therapeutic resistance (e.g., anti PD-1/PD-L1 therapies) (*GeneCards - Human Genes | Gene Database | Gene Search*, n.d.; Melero et al., 2025).

In the cell-membrane fraction of all cells, cell adhesion molecules were detected in high abundance, in particular ITGB1, ITGA6, and CD44 (**Figure 6C**). ITGB1 and ITGA6 form a heterodimer α6β1 that binds laminin and collagen, mediating thus the attachment of cells to the ECM (Su et al., 2024). High expression in cancer cells was associated with stemness and increased metastatic potential (Su et al., 2024). CD44 - a receptor for various ECM components (e.g., hyaluronic acid, MMPs, osteopontin, collagen) that is overexpressed on cancer cells, is a cancer stem cell marker and promoter of metastasis by facilitating adhesion and migration processes (X. Liu et al., 2019; Senbanjo & Chellaiah, 2017). CD44 and ICAM1 (or CD 54, detected in high abundance in HBEC-5i and lower abundance in MDA-MB-231) have been also shown to promote cancer cell clustering ability along with adhesion to endothelial capillary structures (X. Liu et al., 2019; Taftaf et al., 2021).

As expected, the HBEC-5i cells expressed much higher levels of VCAM1 (vascular cell adhesion molecule, or CD106) - an adhesion molecule involved in the reorganization of endothelial tight junctions and mediation of immune cell infiltration from the bloodstream (H. Zhang et al., 2024), and CDH2 (N-Cadherin or CD325) - a molecule that mediates cell-cell adhesion contributing to the stabilization of adherens junctions and integrity of the vascular barrier, as well as to angiogenesis and tumor progression (Kruse et al., 2019; Yu et al., 2019). The cancer cells expressed L1CAM (CD171) and EPCAM (CD326), which are prognostic/diagnostic biomarkers often overexpressed on cancer cells, with roles in supporting cell motility, invasion, proliferation, and metastatic progression (Altevogt et al., 2016; Gires et al., 2020). CDH1 (E-cadherin or CD324), a classical cadherin expressed by epithelial cells that prevents cancer cell invasion by maintaining cell-cell adhesions (Pećina-Šlaus, 2003), was identified only in the cancer cells, in very low abundance in MDA-MB-231 and somewhat higher abundance in SK-OV-3 cells. The SK-OV-3 cells expressed, however, a higher level of CDH2, suggesting the presence of a mesenchymal, more aggressive phenotype. The tight junction proteins TJP1/TJP2 (ZO-1/ZO-2) and OCLN were identifiable, but in much lower abundance than the other cell-membrane proteins, and CLDN5 was not detectable by MS.

### Cell viability and long-term stability of the model system

The endothelial/cancer cell models were maintained for up to 30 days, and cell viability was assessed with red propidium iodide stain in HBEC-5i/SK-OV-3 and a green dead cell stain in HBEC-5i/MDA-MB-231 co-cultures (**Figure 7**). Fluorescence imaging revealed the presence of rather few dead cancer cells on the top of mostly live HBEC-5i cells, indicating that the culture environment and the cancer-endothelial interactions contributed to the survival of cells, the stability of the cellular structures, and the creation of a robust and biologically relevant 3D model of natural tissue.

**Figure 7.**
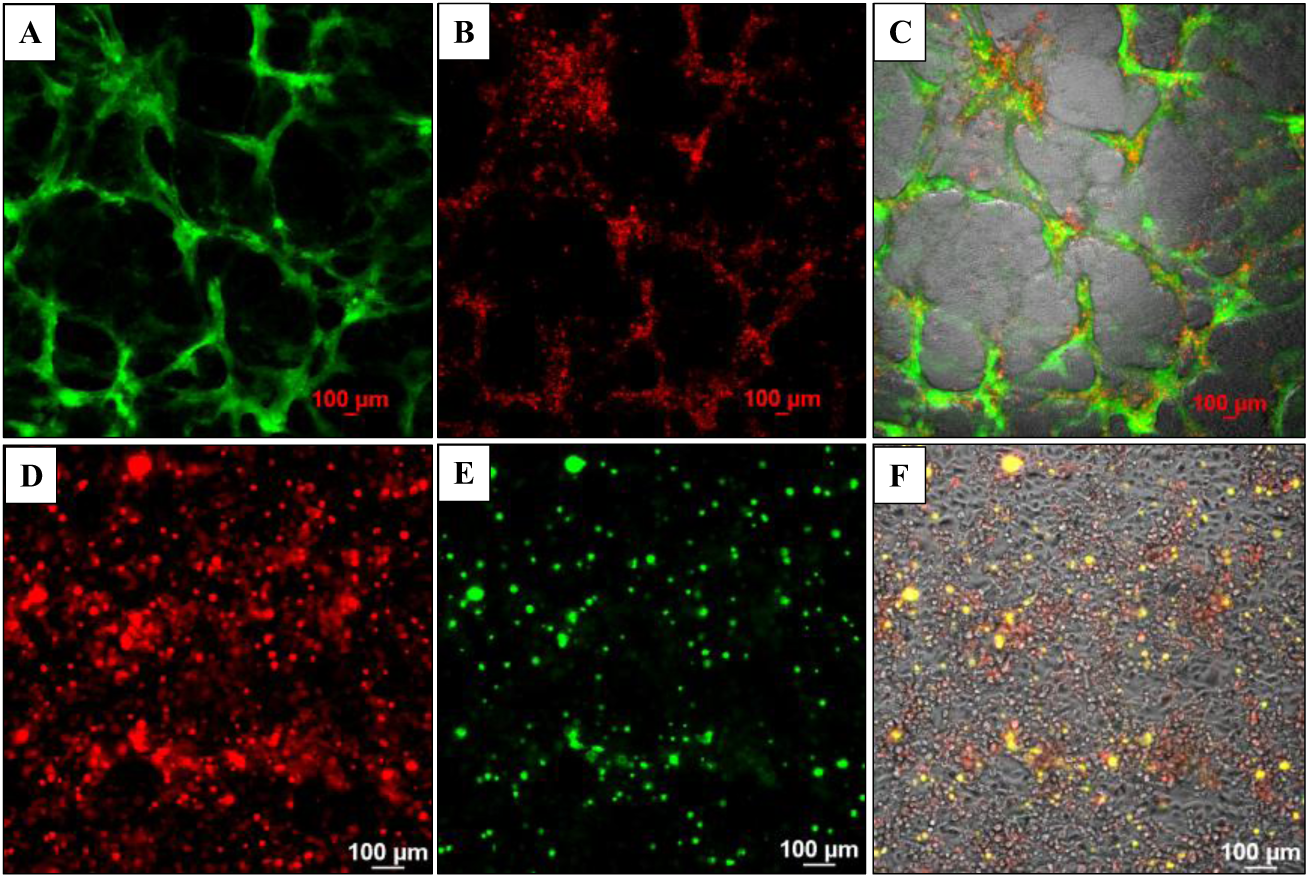
Long-term viability of cells in the pre-mixed endothelial/cancer (1:1) co-culture models assessed by fluorescence microscopy and dead cell staining. Top panel: HBEC-5i/SK-OV-3 (turbo GFP/green) model; (**A**) Green fluorescent SK-OV-3 cells (FITC filter); (**B**) Dead cells (red) visualized by PI staining (TRITC filter); (**C**) Merged image of transmitted, FITC-and TRITC-filtered images of co-cultured cells. Bottom panel: HBEC-5i/MDA-MB-231 (turbo FP602/red) model; (**D**) Red fluorescent MDA-MB-231 cells (TRITC filter); (**E**) Dead cells (green) visualized by staining with EasyProbe dye (FITC filter); (**F**) Merged image of transmitted, TRITC-and FITC-filtered images of co-cultured cells. Images were acquired after 28 days of co-culture.

### Application of 3D co-culture model

As 3D co-culture systems replicate more accurately the *in vivo* environment and enable more reliable predictions of drug efficacy and resistance, the above-described model was evaluated for its suitability in anti-cancer drug screening applications. SK-BR-3 breast cancer cells, which overexpress HER2 receptors and can therefore be effectively targeted with drugs, were used for this purpose. HBEC-5i and red-fluorescent SK-BR-3 cells were mixed in a 1:1 ratio and co-cultured in regular DMEM/HG culture medium until the accumulation of SK-BR-3 cells in the endothelial structures was observed. The constructs presented similar morphological characteristics to the ones observed in the HBEC-5i/MDA-MB-231 model, with cancer cells clustered on the structures created by the endothelial cells. We investigated the effect of Lapatinib, a highly selective, reversible Tyr kinase inhibitor that targets the EGFR (HER1) and HER2 receptors (Opdam et al., 2012), and which possesses the important property of being able to cross the BBB for therapeutic purposes. We have previously shown that the drug can exert broad effects on the SK-BR-3 cells, inhibiting cell-cycle proliferation, adhesion, and migration processes, and leading to cell death and detachment upon prolonged exposure to the drug (Karcini & Lazar, 2022). Here, we compared the drug response of SK-BR-3 cells in 2D monoculture and 3D co-culture systems by visualizing the dead cells after exposure to the drug (**Figure 8**). Confluent monocultures of HBEC-5i and SK-BR-3 were used as controls. There was no impact of drug addition to the HBEC-5i cells, which displayed no increase in the numbers of dead cell counts even after 72 h exposure to Lapatinib (10 µM) (note the few cells labeled with green dead cell stain in **Figures 8A,B**). The SK-BR-3 controls, on the other hand, responded as expected, displaying progressively more dead cells with increased exposure time to the drug, ultimately detaching and leaving large surface areas of the culture plate exposed (note the red/live and yellow/dead SK-BR-3 cells in **Figures 8C,D**, as well as the exposed plate areas). Interestingly, in the 3D co-culture model, the viability of cancer cells was increased, indicating a protective role exerted by the endothelial cells. Major differences between the non-treated co-cultures and the ones treated with 1 µM or 10 µM Lapatinib solution, for 48 h or 72 h, were not observed. Moreover, the cancer cells did not detach from the endothelial cells (**Figures 8E-H**). While more work will be needed to uncover the mechanistic details of this behavior, the results support the hypothesis that the interaction of cancer cells with other cells in the tumor environment increases the resistance to treatment with therapeutic drugs (Salemme et al., 2023).

**Figure 8.**
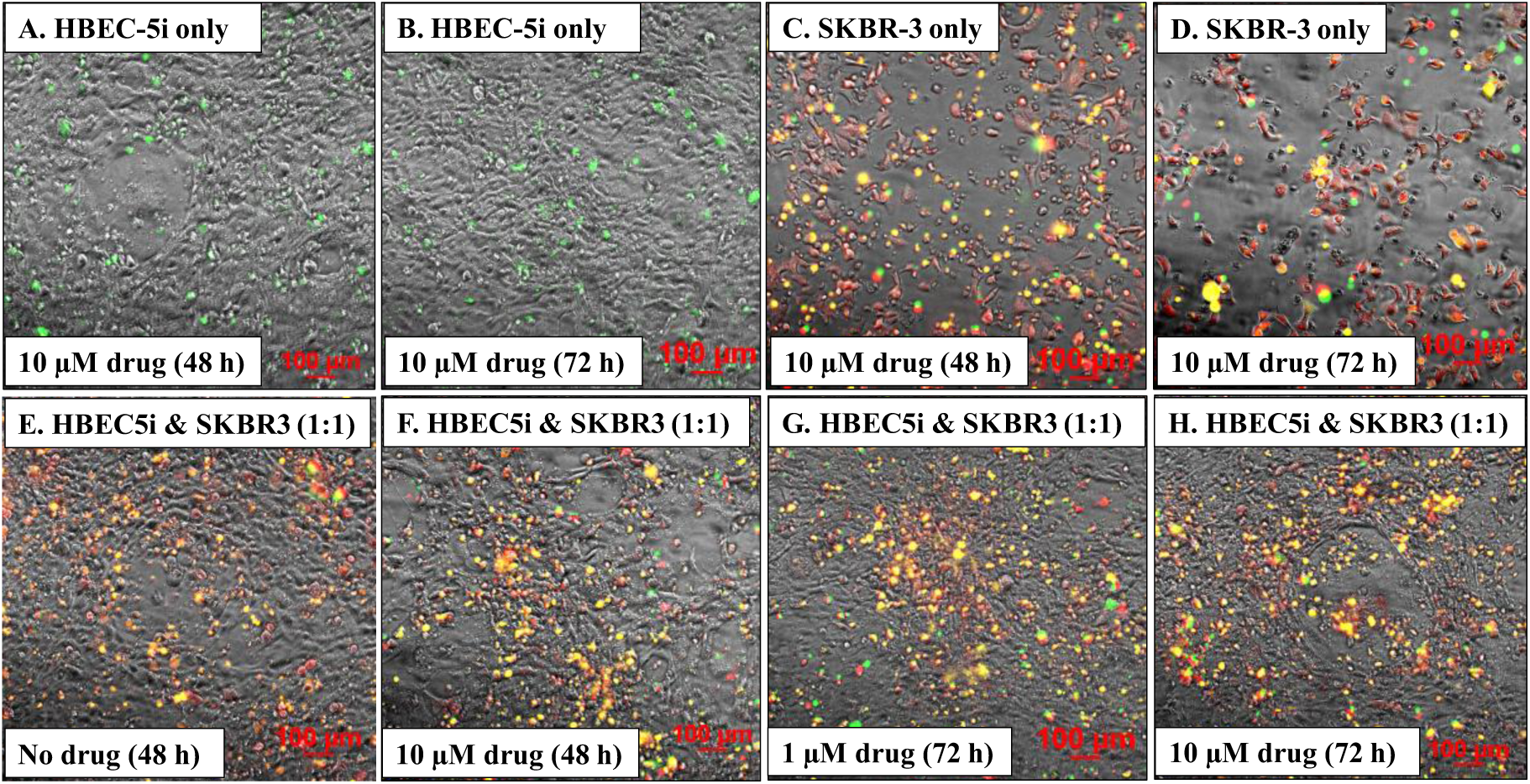
Lapatinib treatment of HBEC-5i and red fluorescent SK-BR-3 cells in mono-and co-culture systems. **Top panel**: mono-culture controls of HBEC-5i and SK-BR-3 cells treated with Lapatinib (10 µM); (**A,B**) Dead cells (green) in HBEC-5i mono-cultures; (**C,D**) Dead cells (yellow) in red fluorescent SK-BR-3 mono-cultures. **Bottom panel**: Co-culture HBEC-5i/SK-BR-3 (turbo FP602/red) cell model treated with Lapatinib (1 and 10 µM). (**E-H**) Dead cells (yellow) in the endothelial/cancer co-culture system. EasyProbe green dye (FITC filter) was used to visualize the dead cells in all cultures.

## Discussion

Intercellular interactions established in the complex TME are essential to cancer cell survival, growth, and metastasis. Due to a limited understanding of these interactions in the unique brain tumor microenvironment, cancer brain metastases are often associated with poor prognosis. While the development of novel *in vivo* mouse models (patient derived xenografts, syngeneic, or genetically engineered mouse models) has significantly advanced our knowledge of tumor biology, the establishment of such models is time consuming and expensive, and the visualization of cancer cell behavior within the TME is challenging. Importantly, interspecies genetic and physiological disparities lead to skewed or inaccurate representations of tumor-host TME interactions, further limiting the use of animal models. Alternative 3D models (e.g., spheroids or organoids) do not fully replicate the complexity of the TME but can simulate key aspects of tumor morphology, composition, and chemical signaling. Nonetheless, difficulties in reproducible culturing and the need for animal-origin or synthetic scaffolding materials that contaminate the environment and induce additional variable effects, impose further restrictions on their use.

In this work we established an *in vitro* scaffold-free 3D co-culture model for studying the interactions between cancer and brain endothelial cells, and their role in tumor development. Abundant collagen production by the HBEC-5i cells created the requisite endogenous substrate for simulating 3D cellular interactions and enabling self-assembly in 3D networked constructs that mimic the natural tissue architecture, in a reproducible and cost-effective manner. The 3D constructs were established without the use of exogenous scaffolding materials or supportive techniques and technologies that help cells self-assemble into aggregates such as physical agitation, hanging drop, liquid overlay, magnetic levitation, bioprinting, or microfluidic guiding (Barbosa et al., 2021; Sharma et al., 2024). While the monoculture of endothelial cells showed the initiation of tube-like, microvascular-mimicking structures after 10 days of culture, in endothelial-cancer co-cultures, these structures were already visible after 6-7 days. The cancer cells affected the phenotype and behavior of the cells in the 3D constructs, as evidenced by the different morphology of the generated constructs. The integrated brain endothelial and highly aggressive cancer cell constructs enabled the examination of cell migration, 3D structure formation, and long-term cell viability, offering valuable insights into the mechanisms that drive tumor cell behavior at the brain metastatic site. The co-culture model was observed up to 30 days, with live-dead cell staining showing only a small proportion of dead cells in the system.

The observed interactions between cancer and endothelial cells were driven by cell-membrane and secreted factors from both cell types. While collagen deposition by HBEC5i cells provided for a fertile matrix that enabled the development of the 3D model, proteomic data revealed that the HBEC-5i, SK-OV-3, and MDA-MB-231 cells expressed a wide array of chemokines and adhesion, growth, and pro-angiogenic factors that supported migration, angiogenic, and proliferation pathways. Chemokine-driven processes in the tumor microenvironment, including signaling sustained by CCL2, CXCL1, CXCL8, CXCL10 and CXCL12, guide not only the migration of both cancer and endothelial cells (Bolitho et al., 2010; Furukawa et al., 2013; Israeli Dangoor et al., 2025; Shen et al., 2022; Zielińska & Katanaev, 2020), but also the remodeling of endothelial cells to ultimately support cancer progression (Salazar & Zabel, 2019). Angiogenic factors secreted by cancer cells (e.g., VEGFs, PDGFs, TGFB1) have been shown to interact with VEGFRs, PDGFRs, Endoglin (ENG), Neuropilin1 (NRP1), and integrin receptors on endothelial cells to activate signaling pathways that promote sprouting angiogenesis and endothelial vasculature in the TME (*GeneCards - Human Genes | Gene Database | Gene Search*, n.d.). Such intercellular interactions that steer the development of microvasculature structures and the migration of cancer cells towards the endothelial cells were observed in the newly developed 3D *in vitro* model system, supporting its physiological relevance. Validation with a transwell assay further confirmed the role of paracrine signaling in shaping the complex tumor microenvironment. Furthermore, the presence of key cell secretome cytokines with roles in innate immunity and inflammatory processes (e.g., CSF1 (*GeneCards - Human Genes | Gene Database | Gene Search*, n.d.)) and various growth factors implicated in the many steps of cancer progression uncovered additional areas of research that could benefit from the model, and prospects for the development of novel prognostic/diagnostic assays and targeted therapies. The co-culture system displayed prolonged stability, holding thus significant potential for investigating anti-cancer drug responses. Subjecting the HBEC-5i/SK-BR-3 3D model to the receptor Tyr kinase inhibitor Lapatinib treatment revealed a differential drug response, characterized by reduced cell morbidity relative to 2D SK-BR-3 monocultures, exposing the protective role conferred by the presence of endothelial cells. These observations underscore the value of the model system as a preclinical research platform for drug testing.

## Conclusions

We developed a novel, scaffold-free, *in vitro* 3D brain endothelial/cancer cell model system that mimics the behavior of cancer cells in the early establishment of the metastatic tumor microenvironment. By using stable transfected fluorescent cancer cells, the model provided a platform for observing the dynamic interplay between cancer and endothelial cells, i.e., the migration, positioning, and attachment of cancer cells to high-density endothelial networked microstructures, and the cancer-cell-induced remodeling of the co-culture environment and morphological changes in endothelial cells. The model presents several unique features when compared to other reported *in vitro* systems. First, the choice of HBEC-5i cells enabled this model to better recapitulate the brain microvasculature due to the expression of well-known tight junction, adhesion, and transport proteins. Second, the model was built based on the inherent high collagen secretion property of HBEC-5i cells without using any commercially available ECM components, thus eliminating contamination risks and making it cost effective. Third, the secretion of key chemokines, pro-angiogenic molecules, growth factors, and ECM enzymes by both cancer and endothelial cells, as well as the expression of specific tight junction and cell adhesion molecules, enabled the spontaneous formation of 3D constructs that mimicked the brain microvasculature with attached cancer cells without any need to control the cell positioning process. Finally, long-term stability and applicability to exploring drug responses underscored the physiological relevance of the model and its potential to accelerate the development of therapeutic interventions. Future work will explore the incorporation of other brain resident stromal and immune cells to better replicate the *in vivo* microenvironment and enable more accurate investigations of the dynamic cellular interactions that delineate the behavior of cancer cells at the brain metastatic site.

## Acknowledgments

This work was supported in part by an award from the National Institute of General Medical Sciences (Grant No. 1R01GM121920) to IML. We thank Yunqian Zhang for establishing the fluorescent MDA-MB-231 cell culture stock. The authors acknowledge the use of Grammarly, Google Gemini, and Open AI ChatGPT for minor editorial improvements to the text, including grammar and sentence syntax. The research conception and design, data acquisition and analysis, and interpretation of results are the sole intellectual contribution of investigators.

## Author contributions

PS performed all experimental work related to the development of the endothelial/cancer co-culture model system, established the fluorescent SK-OV-3 and SK-BR-3 cell culture stocks, and wrote the first draft of the manuscript. SA established the HBEC-5i cell culture stock from ATCC cells and acquired the immunofluorescent microscopy images of HBEC-5i cells. PS and IML conceived the work, and processed and analyzed the data. IML coordinated the work and wrote the final draft of the manuscript. All authors reviewed and approved the final version of the manuscript.

